# Simulating Bead-Tethered Biomolecules: Force-Probe Coupled Steered Molecular Dynamics (fp-SMD) in Optical Tweezers Force Spectroscopy

**DOI:** 10.64898/2026.09.27.754810

**Authors:** Gunan Zhang, Yongsheng Leng

## Abstract

Interpreting single-molecule force spectroscopy (SMFS) data from optical tweezers with steered molecular dynamics (SMD) is hindered by a severe timescale mismatch: SMD trajectories span nanoseconds to microseconds, while SMFS experiments unfold over milliseconds to seconds, forcing SMD to pull six to nine orders of magnitude faster than experiment and systematically distorting rupture forces, molecular pathways, and kinetic intermediates. We introduce force-probe coupled steered molecular dynamics (fp-SMD), which explicitly propagates the force probe (the optically trapped bead) alongside the biomolecule within a single classical Hamiltonian description of the coupled molecule– linker–bead system. A Predict–Correct Trajectory Propagation (PCTP) scheme resolves the resulting computational bottleneck by alternating a bead-propagation step, with the molecular center of mass (COM) held fixed, and a molecular step in which metadynamics and transition-state theory identify the kinetically resolved intermediate reached at each macroscopic time step. On the cholesterol–β-cyclodextrin (β-CD) host–guest complex, fp-SMD reproduces the experimental rupture-force histogram directly at matched conditions, without extrapolation from standard SMD. fp-SMD further resolves how applied force reshapes the molecular escape pathway: we find that cholesterol dissociates via a high-angle route at low force and a direct, small-angle route at high force — providing, to our knowledge, the first direct structural evidence for force-induced pathway switching in a small host–guest system, a mechanism believed to underlie catch-slip bond behavior.

**Significance Statement:** Single-molecule experiments using optical tweezers reveal how mechanical forces unfold proteins, rupture bonds, and drive cell adhesion. Simulations that could explain these experiments at atomic resolution have been hindered by a basic mismatch: they must pull the molecule millions of times faster than a real experiment to be computationally tractable, distorting the forces and pathways they aim to explain. This work introduces a simulation method that explicitly includes the physical probe used in the experiment, not just the molecule, letting simulations match experimental timescales without sacrificing atomic detail. Applied to a cholesterol–cyclodextrin complex relevant to atherosclerosis and drug delivery, the method shows how force redirects a molecule’s escape pathway — a mechanism believed to underlie many force-sensitive switches.

---

Single-molecule force spectroscopy (SMFS) using optical tweezers, magnetic tweezers, or atomic force microscopy (AFM) has become a central tool for probing the mechanics of individual biomolecules and biomolecular complexes under mechanical load[1]. Unlike ensemble-averaged thermodynamic measurements, SMFS resolves force-dependent behavior of individual molecular pairs, giving direct access to rupture forces, bond lifetimes, and — when combined with theory — the underlying kinetic pathways and energy landscapes that govern molecular association and dissociation.

A long-standing and still poorly understood question in mechanobiology is whether, and how, applied mechanical force redirects the pathway by which a biomolecular complex assembles or dissociates. This question sits at the heart of some of the field’s most striking phenomena, including catch-slip bond behavior — the counterintuitive increase, followed by decrease, of bond lifetime with applied force observed in many cell-adhesion molecules[2-4], a signature widely believed to reflect force-induced switching between distinct structural pathways. Recently, catch-slip-like behavior has also been found in small, non-protein host– guest systems (H. Mao, unpublished data), extending this question beyond the classical adhesion-receptor context. Such pathway switching, and hence the transition from catch to slip bonds, has so far been inferred only indirectly from macroscopic force–lifetime curves[2]. Direct evidence for a force-induced switch in the geometric dissociation route itself has been largely inaccessible, in part because few computational tools have been able to resolve molecular dynamics (MD) at both the force regime and the timescale of the experiment. Recent enhanced-sampling MD approaches have begun to close this gap for protein–protein catch bonds[4], but a directional change in the dissociation pathway geometry has not yet been directly observed. Conventional steered molecular dynamics (SMD) has separately resolved force-induced hydrogen-bond formation within a single, non-switching cadherin conformation[3], though at pulling velocities orders of magnitude faster than the experimental catch-bond measurements[5].

While SMFS measures forces and extensions, its computational counterpart, SMD, can in principle resolve the underlying atomistic pathways and energy landscapes. In practice, however, SMD pulls six to nine orders of magnitude faster than SMFS experiments. This timescale gap severely limits direct, quantitative comparison between simulated and measured force spectra, and leaves the question of force-dependent pathway switching largely unresolved.

Here, we introduce force-probe coupled steered molecular dynamics (fp-SMD), a simulation method designed to close this gap by explicitly and self-consistently propagating both the biomolecule and the force probe — the optically trapped bead — under a single, rigorously derived Hamiltonian description of the coupled experimental system. We illustrate and validate the method on the cholesterol–β-cyclodextrin (β-CD) host–guest complex studied by optical tweezers SMFS in our previous work[6], a system of relevance to cyclodextrin-mediated dissolution of pathological cholesterol crystals in atherosclerosis[7, 8].

## Computational Tools for Interpreting SMFS Data

Several classes of computational method have been developed to bridge molecular simulation and SMFS experiment. SMD [9-11] remains the default tool: an external bias pulls a chosen atom or the molecule’s center of mass (COM) along a reaction coordinate, directly paralleling AFM, optical-tweezers, and magnetic-tweezers protocols [1]. Unlike the physical experiment, however, the simulated system has no explicit cantilever, bead, or molecular linker; the force probe’s mass, compliance, and stochastic dynamics are simply absent [12]. This omission is consequential: a growing body of work shows that the physical linker and tethering object used in SMFS measurably shape the inferred kinetics, reducing the effective diffusion coefficient sampled [13] and requiring the probe’s own dynamics to be explicitly deconvolved from the measured signal [14-16]. Existing strategies include Jarzynski-equality free-energy reconstruction [17], Bell–Evans/Kramers rate-theory extrapolation [18-22], enhanced-sampling protocols that extend the accessible force or velocity range at the cost of a continuous trajectory [23-28], matched high-speed AFM/MD studies that push experiment toward simulation timescales [22, 29], and coarse-grained stochastic models that replace the coupled molecule-plus-probe system with a reduced diffusive coordinate [22, 30]. Each addresses only part of the problem; none explicitly and self-consistently couples the atomistic dynamics of the biomolecule to the physical dynamics of the force probe used in the actual optical-tweezers experiment. Without such coupling, the timescale mismatch discussed below cannot be treated on the same footing as the experiment.

## The Timescale Problem in SMD

Optical tweezers operate with instrument time resolution of 0.1–10 ms over pulling experiments spanning milliseconds to seconds[1, 15], while routine all-atom SMD is limited to nanosecond–microsecond trajectories[12]. To observe rupture within this shorter simulation window, SMD must use loading rates of 10^8^– 10^11^ pN/s — six to nine orders of magnitude faster than experiment. This mismatch has three consequences: (1) systematic overestimation of rupture forces, since pulling faster than the molecule’s intrinsic relaxation drives the system out of quasi-equilibrium; (2) distortion or masking of true kinetic pathways, forcing the system through a single, non-physiological dissociation route; and (3) loss of a well-defined bound-state lifetime, since lifetime-versus-force curves computed at artificially fast pulling speeds do not correspond to any physically realizable timescale. Extrapolation schemes based on the Bell–Evans model[18, 19] can partially correct simple, single-barrier systems[20, 21], but cannot be relied on once additional barriers or intermediates become relevant as the loading rate[22] or force[2, 3, 5] changes, which is exactly the regime of interest studied here. Closing this gap requires a method that keeps the biomolecule in near-equilibrium, physically meaningful conditions while still reaching the macroscopic timescale of the force probe. fp-SMD meets this requirement by propagating the probe explicitly together with the molecule, then using PCTP to advance the probe in macroscopic time steps while the kinetically resolved molecular state is updated at each step.

## 1. Computational Approach

### Model System

We use the cholesterol–β-cyclodextrin (β-CD) host–guest complex as the model system to develop and illustrate fp-SMD. This model system was studied experimentally using optical tweezers and computationally using standard SMD in our previous work[6]. The schematic of the molecule–linker–bead system is shown in **Fig. 1**. Cholesterol is pulled from its hydroxyl end relative to a fixed β-CD (only the central glucopyranose ring of β-CD is held fixed). In the experiment, mechanical pulling is achieved by moving one optical trap at constant velocity *V* (force-ramp pulling) relative to the fixed β-CD.

**Fig. 1.**
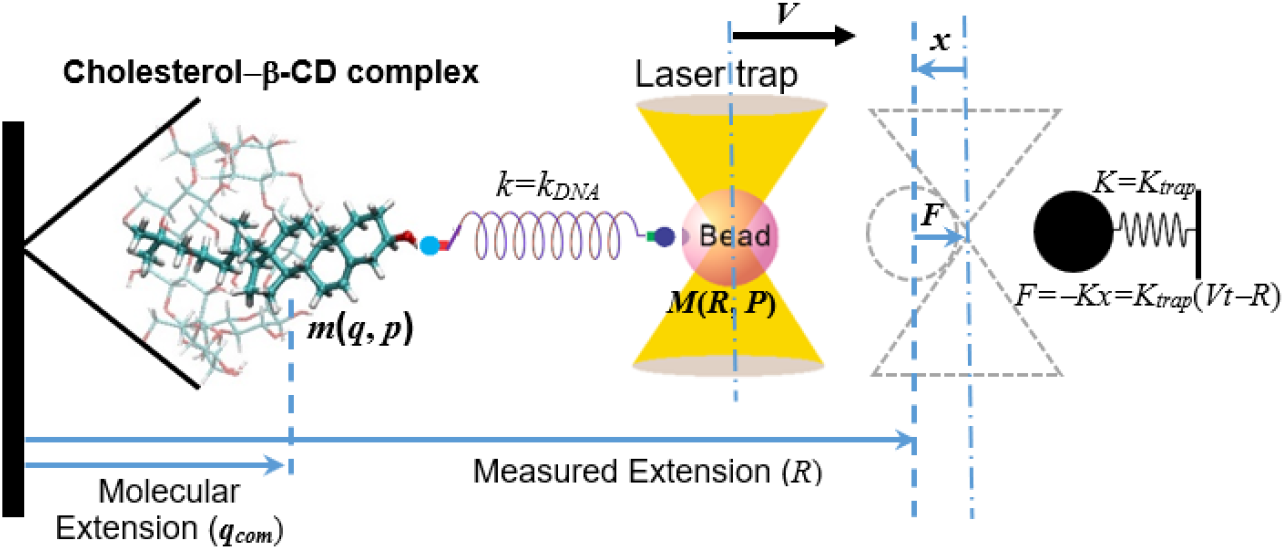
Mechanical model of cholesterol−β-CD host-guest complex in optical tweezers.

Two independent, experimentally and computationally relevant coordinates describe the system, both referenced to an arbitrary zero plane: *R* — the measured extension of the molecule–linker–bead system (the bead position), and *q*_com_ — the center-of-mass position of the cholesterol molecule, which is *not* directly observable experimentally[15].

If the bead has mass *M* and the optical trap has restoring force constant *K*_*trap*_, the only experimentally measurable force in optical tweezers is the trap restoring force,

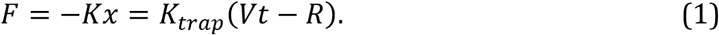

### Hamiltonian Formulation

fp-SMD is built on a rigorous classical Hamiltonian treatment of the combined molecule–linker–bead system, extending our previously developed theoretical framework for AFM-based force spectroscopy of gold nanojunctions[31] to soft, solvated biomolecular systems.

The free energy of the combined system is

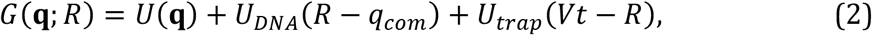

where *U*(**q**) is the potential energy of the host–guest molecular complex (the coordinates **q** representing all explicit atoms of cholesterol and β-CD), *U*_*DNA*_(*R* − *q*_*com*_) is the potential of the DNA linker connecting the molecule to the bead, and *U*_*trap*_(*VV* − *R*) is the potential energy of the optically trapped bead.

The total Hamiltonian of the coupled system is *G*(**q**; *R*) plus the kinetic energy terms of both molecule and bead:

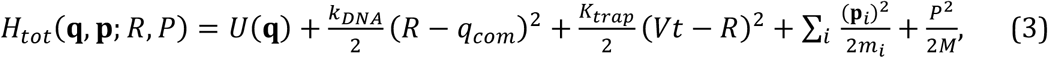

where *m*_*i*_ and **p**_*i*_ are the mass and momentum of atom *i* in the host–guest complex, and *P* is the momentum of the bead. Because the DNA linker’s mass is negligible compared to the bead, only its harmonic potential *U*_*DNA*_ = *k*_*DNA*_2 (*R* − *q*_*com*_)^2^ is retained, with the effective spring constant *k*_*DNA*_ obtained from the worm-like chain (WLC) model of DNA elasticity[32, 33].

### Coupled Equations of Motion

Trajectory propagation based on the total Hamiltonian (3) includes solving two sets of equations of motion[34]: (i) 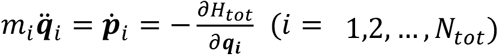 for all-atom MD trajectory propagations of the cholesterol−β-CD molecular complex that has *N*_*tot*_ atoms, and (ii) 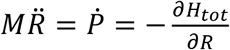 for the bead trajectory propagation. Using Eq. (3), these two sets of equations of motion can be rearranged as

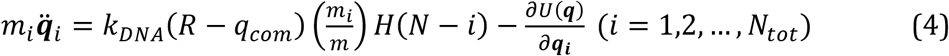

for each atom in the molecular complex, and

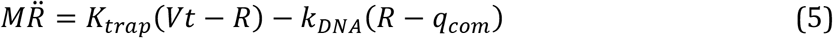

for the bead.

In Eq. (4), *m* and *N* are the total mass and number of atoms of the cholesterol molecule. *N*_tot_ − *N* accounts for the remaining atoms in β-CD. *H*(*n*) is a Heaviside function (*H*(*n*) = 1 for *n* ≥ 0 and *H*(*n*) = 0 for *n* < 0) to distinguish between cholesterol and fixed β-CD. Summing over all individual equations in (4) up to *N* atoms in cholesterol and noting that 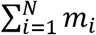 — the dynamic inertia force of the cholesterol, we obtain the equation of motion for the cholesterol apart from β-CD as

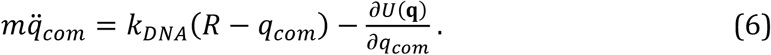

Now if the cholesterol molecule is always in near-equilibrium condition, as we assume in SMFS experiment, then the DNA linker force (see **Fig. 1**) is a direct measure of the *intermolecular mechanical force (IMMF)* [6] between the cholesterol and β-CD in the host-guest complex, i.e.,

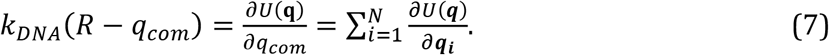

However, as we discussed previously, the only *measurable* force in optical tweezers is the restoring force, *F = K*_*trap*_ (*Vt* − *R*), through the measured extension *R*.

Given that the SMFS experiment is carried out in aqueous media, the Brownian motion of the bead should be considered according to the fluctuation-dissipation theorem. This is achieved by adding two additional stochastic forces terms to the bead Eq. (5): the hydrodynamic drag force 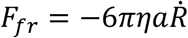 (with *π* the dynamic viscosity of water and *π* the bead radius), and the random thermal force *F*_*th*_, with ⟨*F*_*th*_(*V*)⟩ = 0 and autocorrelation ⟨*F*_*th*_(*V*)*F*_*th*_(*V*′)⟩ = 12*πππk*_*B*_ *T δ*(*V* − *V*′). The final coupled equations of motion governing the host-guest system are given by Eq. (6) for the cholesterol molecule and Eq. (8) for the optical-tweezers bead:

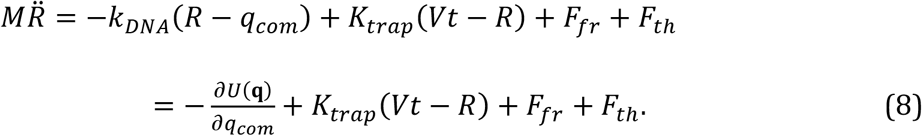

The second form of Eq. (8) follows from Eq. (7); Eq. (4) describes all individual atoms of the cholesterol/β-CD complex explicitly, while Eq. (6) simultaneously propagates the cholesterol COM. *SI Section S1* validates that Eq. (8) correctly reproduces the Brownian motion of a bead in a harmonic optical trap against experimental optical-trapping interferometry data[35]. In standard SMD, the molecular potential also serves as the potential of mean force (PMF) along the pulling coordinate, whose derivative defines the true molecular force; in fp-SMD, the trap restoring force in Eq. (8) instead corresponds directly to the measured IMMF, with the difference accounting for hydrodynamic drag, thermal noise, and bead inertia force. Consequently, whereas real SMFS experiments can recover the true PMF only through a nontrivial deconvolution procedure[16], fp-SMD provides direct access to this quantity.

### The Timescale Bottleneck in Direct Propagation

Equations (6) and (8) fully define the fp-SMD model, but they cannot be integrated directly and efficiently: because the cholesterol molecule and the optically trapped bead operate on vastly different characteristic timescales, simultaneously tracking both sets of degrees of freedom at the atomistic integration step needed for Eq. (6) is computationally prohibitive over the macroscopic time span needed for Eq. (8). Resolving this bottleneck — while preserving the physical coupling between molecule and bead — is the purpose of the Predict–Correct Trajectory Propagation (PCTP) scheme introduced below.

### The PCTP Algorithm

The central idea underlying PCTP is that the molecule and the bead can be advanced in alternating stages without breaking their physical coupling: the bead equation (Eq. 8) is propagated using macroscopic time steps, while the molecular equation (Eq. 6) (also Eq. 4) is solved separately, at each macroscopic step, to find the kinetically-resolved intermediate state of the molecule for the current bead position. Agreement between the resulting fp-SMD-computed IMMF and the experimentally measured IMMF supports this alternating scheme.

The following first-order analysis illustrates the mechanism of the PCTP integration scheme in fp-SMD. In Eq. (8), the activation force, *F*_*a*_(***q***, *R*)= − *k*_*DNA*_(*R* − *q*_*com*_) + *K*_*trap*_(*Vt* −*R*), drives the system through a nonequilibrium pulling process. Suppose that at time *t*, the molecular conformation of the host-guest complex is ***q***(*t*) and the measured extension is *R*(*t*) (see **Fig. 1**), yielding an activation force of *F*_*a*_(***q***(*t*), *R*(*t*)). When the laser trap moves by *VΔt*, the measured extension *R* increases by a small increment *ΔR*. Simultaneously, the molecular conformation shifts to ***q***(*t*)+Δ***q***, updating the COM position of the cholesterol to *q*_*com*_(*t*) + Δ*q*_*com*_, and the activation force to *F*_*a*_(*t*+Δ*t*) = *F*_*a*_(***q***+Δ***q***, *R*+Δ*R*). To the first-order approximation, this updated force expands as: *F*_*a*_(***q***+Δ***q***, *R*+Δ*R*) ≈ *F*_*a*_(***q***, *R*) + (∂*F*_*a*_/∂***q***)Δ***q*** + (∂*F*_*a*_/∂*R*)Δ*R*. Because the ***q***-dependent and *R*-dependent contributions to *F*_*a*_ enter this expansion linearly, they can be evaluated in separate stages rather than solved jointly, motivating the Prediction–Relaxation–Correction sequence below. Within each step, however, the corresponding contribution is evaluated exactly rather than truncated at first order.

Figure 2 summarizes the three-step PCTP cycle. In **Step 1**, the cholesterol COM is held fixed at its previous-cycle thermodynamic average while Eq. (8) is integrated over the macroscopic step ΔT. Because the activation force is harmonic in *R* at fixed *q*_com_, this update is exact rather than a first-order truncation. The only approximation is that *q*_com_ can be held fixed over ΔT, i.e., that its excursion stays within ∼1 nm (see Remark 1; the safe increment *ΔR′* is set by the linear-response regime, *SI Section S2*). In **Step 2**, using the updated *R* as a fixed constraint, the host–guest complex is relaxed via well-tempered metadynamics[27] (implemented via the PLUMED plugin[36, 37]) to map the free-energy surface and PMF along the pulling pathway; transition-state theory converts the resulting barriers into a rate matrix **K**[38-40], and propagating the master equation to the actual cycle time ΔT identifies the kinetically resolved intermediate the system actually reaches within ΔT, rather than assuming instantaneous equilibration. We use a single, state-independent rate pre-factor k_0_ applied uniformly across **K**, with only the barrier ΔG varying by pair (*SI Section S3*). In **Step 3**, the updated molecular coordinate is used to re-evaluate the molecular force and solve Eq. (8) again, correcting the predicted extension *R* + Δ*R′* to the final *R* + *ΔR*; the resulting IMMF is compared against the optical-tweezers measured value. Steps 2 and 3 are repeated iteratively until the change in bead position between iterations falls below a preset convergence threshold (∼1 nm, smaller than the bead’s typical thermal fluctuations, Fig. S2); in practice this inner loop typically converges within a single iteration. This scheme couples molecular-level sampling to the macroscopic integration of force-probe motion, providing a computational framework for force-driven biomolecular processes that reaches timescales far closer to biologically relevant, slow motions and fluctuations underlying real mechanobiological function than standard SMD permits.

### Simulation Parameters and Package

Preliminary fp-SMD/PCTP simulations of the cholesterol–β-CD system are based on our previously published work[6]. The CHARMM36m force field[41] is used throughout. The fp-SMD/PCTP package is implemented as an external wrapper calling the LAMMPS MD engine[42, 43]. SMFS optical tweezers parameters are set to: bead mass *M* = 5 × 10^−15^ kg, optical trap stiffness *K*_*trap*_ = 132.4 µN/m, and DNA linker stiffness *k*_*DNA*_ = 650 µN/m. A loading rate of 5.5 pN/s was used, equivalent to a pulling speed of *V* = 50 nm/s at an effective stiffness of 110 µN/m (= *Kk*/(*K* + *k*)), matching the slope of the experimental SMFS force–extension curve in the 10–30 pN range[6]. Each PCTP cycle integrated over a macroscopic time *ΔT* = 20 − 100 ms (internal step *ΔV* = 200 ns, 100,000 - 500,000 steps per cycle), producing a 10 − 50 Å pulling distance per cycle, which is inside the thermal fluctuations (±80–90 Å quadratic region established from Fig. S6). Because the molecular conformation ***q*** and cholesterol center-of-mass *q*_*com*_, thus the molecular force term 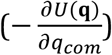 in Eq. (8), is updated once per PCTP cycle, *ΔT* itself — not the internal integration step *ΔV* — sets the effective temporal resolution of coupled bead–molecule dynamics in fp-SMD. This resolution (20 − 100 ms) is comparable to / coarser than typical time resolution of optical tweezers instruments (0.1–10 ms)[1]; refining *ΔT* toward instrument resolution is a natural target for future development. We benchmark the PCTP framework by evaluating its ability to predict rupture forces and capture unbinding escape pathways. All fp-SMD/PCTP simulations are performed at T = 296 K.

**Fig. 2.**
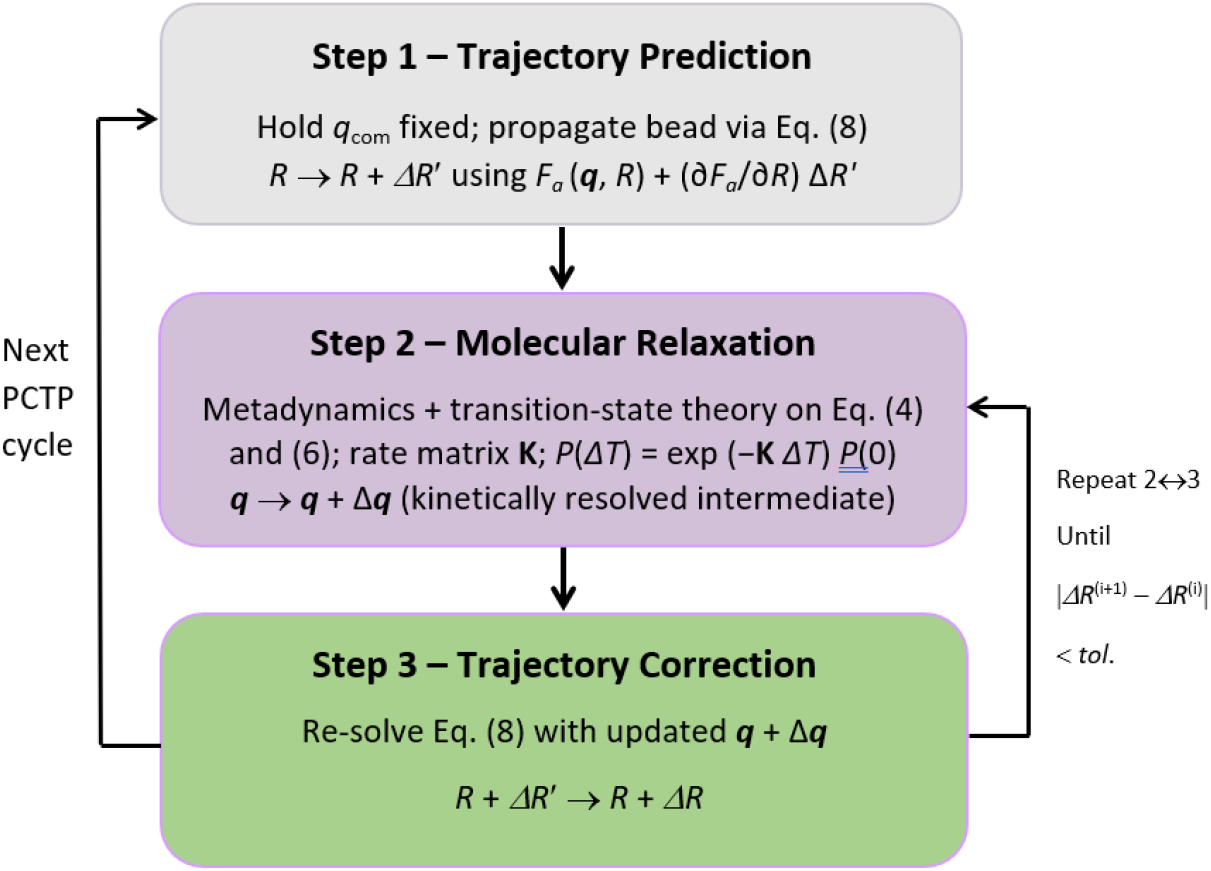
The three-step PCTP prediction-relaxation-correction cycle. Steps 2 and 3 are repeated until convergence before the next macroscopic cycle begins.

## 2. Results and Discussion

### Rupture-Force Validation

We validate fp-SMD/PCTP against the cholesterol–β-CD complex pulled from the hydroxyl end, the one geometry for which both optical-tweezers measurements and standard SMD simulations are directly available for comparison[6]. Experimentally, pulling the host–guest complex from the hydroxyl head group of cholesterol at a loading rate of 5.5 pN/s (171 independent runs) yields a single rupture-force population centered at 32 pN, with a corresponding change-in-contour-length of 8.1 nm. Standard nonequilibrium SMD simulations of the same pulling geometry performed at a pulling speed of 0.2 m/s (25 independent runs), likewise yield a single rupture-force population, but centered at 136 pN: roughly 4.3 times higher than the experimental value[6] as shown in **Fig. 3** (top). Both experimental measurement and SMD simulation report a single dominant population for this pulling direction, consistent with the free-energy landscape found for hydroxyl-end pulling — a primary minimum with only a shallow secondary basin offset by about 2.0 Å, with a barrier of order 1 k_B_T (see the original Fig. 3B in [6]). This is therefore a clean, direct comparison, not complicated by the multi-population behavior seen for alkyl-end pulling[6]. The ∼four-fold discrepancy is the system-specific signature of the timescale/loading-rate mismatch discussed in the timescale problem in SMD. The standard SMD pulling speed of 0.2 m/s with a spring constant of 2 kcal/(mol Å2) (= 1.39 N/m)[6] gives a loading rate of 2.8 × 10^11^ pN/s, about ten orders of magnitude higher than the loading rate used experimentally (5.5 pN/s). We find that this loading rate difference likely accounts for most of the four-fold shift in the observed rupture force.

**Fig. 3.**
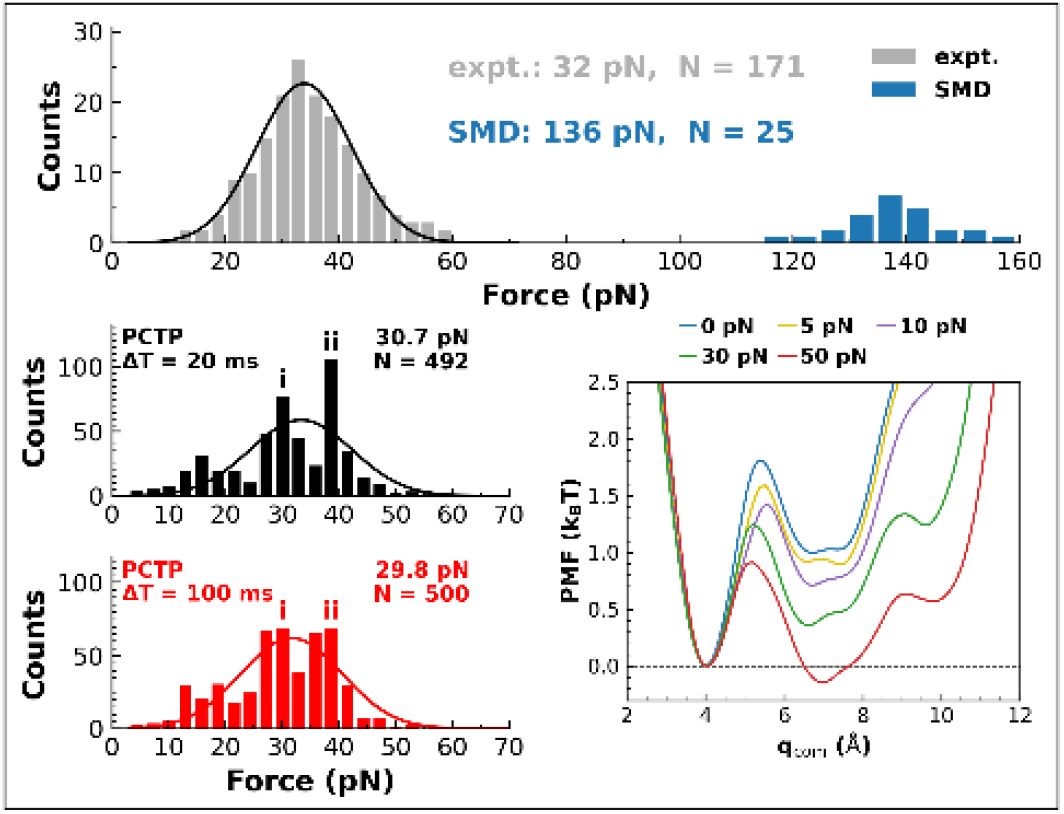
Comparison of force histograms obtained from fp-SMD/PCTP simulations and SMFS experiment in optical tweezers. *Top:* raw experimental (32 pN, N = 171) and standard SMD (136 pN, N = 25, pulling speed 0.2 m/s) force histograms, adapted from our previous work.^6^ *Bottom left and center:* fp-SMD/PCTP force histograms obtained using the force-controlled protocol (SI Section S5) at two PCTP macroscopic time steps, ΔT = 20 ms (N = 492, mean force 30.7 pN) and ΔT = 100 ms (N = 500, mean force 29.8 pN); each histogram shows two force peaks at 30 pN (peak i) and 39 pN (peak ii), absent from the raw experimental histogram. *Bottom right:* PMF profiles of cholesterol along the pulling coordinate at five representative loads (0, 5, 10, 30, and 50 pN), showing a shallow secondary free-energy minimum (offset by 2.5–3 Å from the primary minimum, with a barrier of 1–1.9 k_B_T).

By contrast, fp-SMD/PCTP simulations for the same pulling geometry are carried out at the experimental loading rate of 5.5 pN/s, equivalent to a pulling speed of 50 nm/s at the effective trap–linker stiffness (110 µN/m) used here. Using well-tempered metadynamics sampling[27] in PCTP Step 2, all the free-energy surfaces (FESs) and the PMF profiles, together with all sampled conformations under different loads, are precalculated and stored in a large dataset for extensive fp-SMD/PCTP runs (*SI Section S4*). These conformations, because they are still under artificial Gaussians, are then de-biased using short (∼2 ps) unbiased MD runs to transition to *kinetically resolved intermediates*[44, 45]. Application of the pre-calculated FESs and PMF profiles in fp-SMD/PCTP runs is detailed in *SI Section S5*, where a force-controlled protocol is adopted in the force-ramp pulling.

Figure 3 shows that fp-SMD/PCTP reproduces the experimentally measured rupture-force distribution directly, without extrapolation or rescaling. Here we show two simulation results at two PCTP macroscopic time steps: ΔT = 20 ms, N = 492, F = 30.7 pN; and ΔT = 100 ms, N = 500, F = 29.8 pN. Both mean forces closely match the experimental value (32 pN, N = 171), and, despite a five-fold difference in ΔT, the two distributions are consistent with each other. Surprisingly, each fp-SMD/PCTP distribution reveals two distinct force peaks, at 30 pN (peak i) and 39 pN (peak ii). These peaks are absent in the raw optical-tweezers histogram. We speculate that the raw histogram has not been deconvolved, as was done for the alkyl-end data[6]. Temporal resolution is unlikely to be the cause: the simulated peaks are insensitive to ΔT (20 vs. 100 ms), and the experimental time resolution (∼10 ms) is finer than or comparable to the PCTP macroscopic time steps used here. Deconvolving the existing experimental data would therefore provide a direct test of the predicted two-peak structure. The two peaks are consistent with the shallow secondary free-energy minimum previously identified for this pulling direction[6]. Furthermore, our current fp-SMD/PCTP simulations for hydroxyl-end pulling directly reproduce this detailed PMF profile, which is evidently force-dependent, a key property our previous work using umbrella sampling does not directly provide[6]. As shown in the bottom right in **Fig. 3**, the second basin is separated from the primary minimum toward the β-CD exit by 2.5–3 Å, with an energy barrier of 1–1.9 k_B_T (see also Fig. S7(C)).

### Force-Dependent Escape Pathway Switching

Beyond reproducing the rupture forces measured by optical tweezers SMFS, fp-SMD resolves how applied force reshapes the molecular escape pathway. As shown in **Fig. 4**, at low force (5 pN), cholesterol escapes β-CD via a high-angle pathway; at high force (50 pN), it instead escapes via a direct, small-angle pathway. This pathway shifting is also seen in the 2D-FES evolutions in Fig. S7(A). To our knowledge, this is the first direct structural demonstration that mechanical force can redirect the geometric route of a biomolecular dissociation pathway at the correct experimental timescale—the force-dependent pathway switching believed to underlie catch-slip bond behavior[2, 3]. Recent enhanced-sampling MD studies have begun to simulate force-dependent unbinding kinetics and the intrinsic, contact-based catch-bond behavior of individual bound states in the vinculin–actin system[4]. These simulations reveal that the escape geometry differs between bound states and between pulling directions, but a force-induced switch in the dissociation pathway of a given bound state has not, to our knowledge, been directly resolved.

**Fig. 4.**
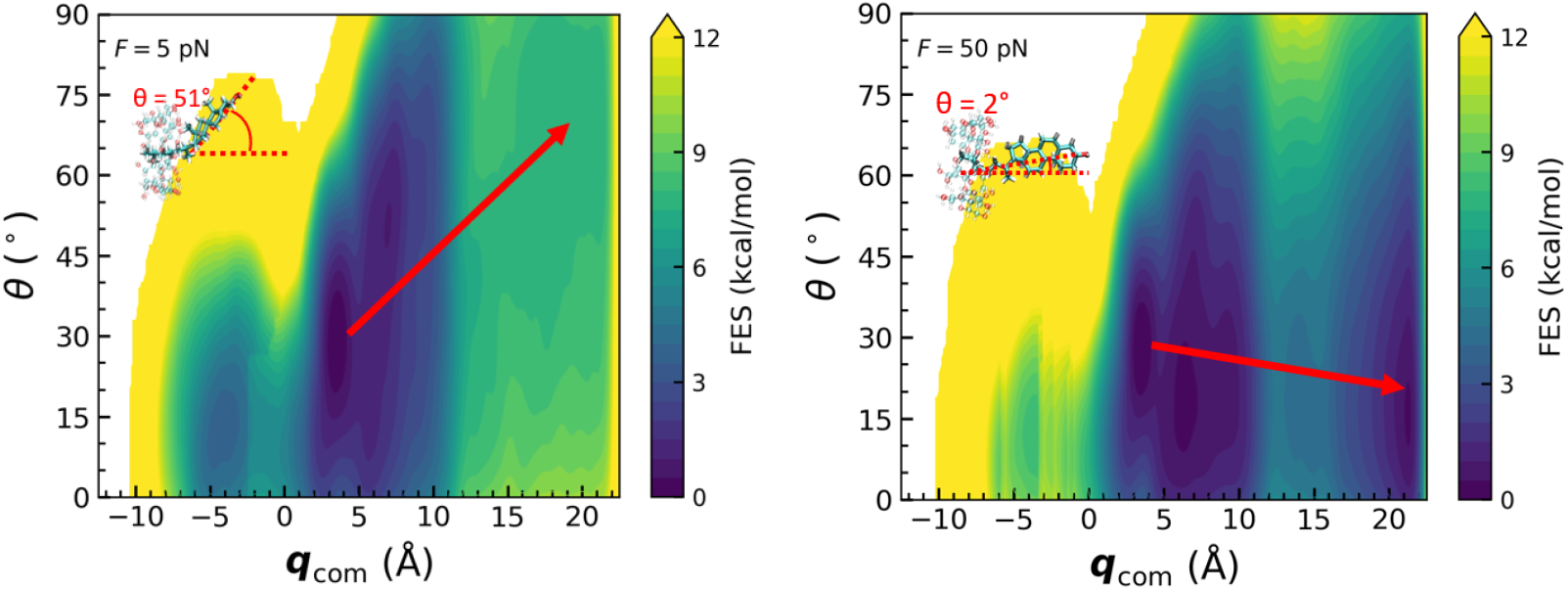
Free energy surface (FES) of cholesterol in β-CD under different external loads: (A) F = 5 pN; (B) F = 50 pN. The tilt angle (θ) and the center-of-mass position (*q*_com_) of the cholesterol molecule are selected as the two collective variables (CVs) in metadynamics sampling. The red arrow indicates the direction of cholesterol exiting the β-CD.

This pathway dependence has a clear physical origin, evident from PCTP’s molecular relaxation trajectories. In pure molecular relaxation runs (bead held static), cholesterol near the β-CD exit shows large-amplitude tilt-angle swings, occasionally drawing the molecule back into the pocket (due to strong hydrophobic interactions between cholesterol and β-CD) before full escape; once fully seated within β-CD, however, its conformation is comparatively rigid, with only small-amplitude fluctuations. This asymmetry explains the two pathways observed: at low force, cholesterol has time to explore these tilt-angle swings before exiting, favoring the high-angle route; at high force, it is pulled through too quickly for such swings to develop, favoring the direct, small-angle route. A detailed conformational analysis of all independent PCTP runs shows that cholesterol — right before its exit from β-CD — largely adopts the high-angle or direct, small-angle exit conformation under low- and high-force rupture, respectively (**Fig. 4**). This observation also validates the tilt angle (θ) and cholesterol center-of-mass (*q*_*com*_) as the two collective variables used in metadynamics sampling, since the conformational flexibility driving pathway selection is concentrated in the exit region rather than in the stable, fully encapsulated state.

These force-dependent escape pathways offer a mechanistic basis for how mechanical conditions — such as shear forces encountered during cholesterol removal from membranes or arterial plaques — may influence the efficiency of cyclodextrin-mediated extraction of cholesterol. A capability-by-capability comparison between standard SMD and fp-SMD/PCTP, spanning physical model, trajectory continuity, timescale accessibility, loading-rate realism, and pathway/lifetime resolution, is summarized in *Table S1 in SI Section S6*. Three remarks need to be further discussed:

#### (1) Molecule center-of-mass (COM) frozen approximation

In the Prediction step (Step 1), PCTP takes the molecular COM position (q_com_) at its previous-cycle thermodynamic average as a fixed value while the bead is propagated through ΔT according to Eq. (8), which depends on the molecule only through this q_com_. Here, thermal fluctuations of the COM within a basin (≈ 0.02–0.03 nm, SI Figs. S3 and S4(b)) average out over the bead’s response time (τ_A_ ≈ 24.7 μs, SI Section S1), so the bead responds only to the mean q_com_. Beyond these small thermal fluctuations, the frozen-COM approximation can be further relaxed, as long as the COM displacement (δq_com_) during ΔT remains small enough that the force transmitted to the molecule is essentially unchanged. Because the bead follows the COM through the stiff DNA linker, we show that for a 1 nm COM displacement, the force change transmitted to the molecule is only about 0.11 pN (SI Section S7). We take ∼1 nm as the reference displacement, equal to the transition-state distance x^‡^ ≈ 10 Å (SI Fig. S7(C)). In the extreme case where the molecular COM moves across the full range of the PMF (≈ 1.8 nm, SI Fig. S7(C)), we find that the force changes by only ≈ 0.2 pN. Displacements beyond x^‡^ correspond to barrier crossing. In this case, Step 2 identifies the new molecular state from the precalculated large dataset of molecular meta-conformations, the updated COM is then taken from this identified meta-conformation (SI Section S5), and Step 3 solves Eq. (8) again with this new q_com_. Therefore, PCTP does not require the molecule to relax faster than the bead moves. Instead, PCTP relies only on the standard thermally activated, transition-state-theory description of barrier crossing, as in Tiwary and Parrinello[44, 45]. We expect the frozen-COM approximation to be challenged only by much larger conformational rearrangements, stiffer DNA linkers, or much longer ΔT, and mapping these regimes is left for future work.

#### (2) Collective-variable selection

The molecular relaxation step relies on metadynamics sampling along a small set of collective variables (here, the tilt angle *θ* and *q*_com_). While these CVs were validated post hoc by the physical picture in **Fig. 4**, poor CV choice in general risks missing hidden slow degrees of freedom, particularly in larger or more conformationally complex systems. Extending PCTP to such systems, such as cholesterol monohydrate bundles, larger biomolecules (proteins, nucleic acid structures), and multi-component aggregates, will require testing free-energy landscape roughness, CV adequacy and convergence, and the generality of the frozen-COM and transition-state-theory assumptions underlying PCTP, a natural next step building directly on the present benchmark.

#### (3) Catch-slip curves and transition path times

The present results show force-dependent pathway switching at two force values (5 and 50 pN), but do not yet give a full catch-slip bond–lifetime curve. We expect that such a curve can be obtained from PCTP by repeating the transition-state-theory analysis over a range of forces. Transition path times can in principle also be obtained. In optical tweezers experiments, the transit time is the time that the measured bead extension stays between the two boundaries of the barrier region[46, 47]. In PCTP, this time follows from Eq. (8): we propagate the bead with the molecular COM updated at each cycle, and accumulate the time the bead spends inside the barrier region. Because the stiff DNA linker makes the bead follow the molecular COM (SI Section S7), the bead relaxes toward the COM with time constant τ_A_ ≈ 25 μs (SI Section S1). If the molecular crossing is much longer than τ_A_, the bead simply trails the COM with a constant delay, which shifts the entry into and exit from the barrier region equally, so the bead transit time equals the molecular crossing time. If the crossing is comparable to or shorter than τ_A_, the bead needs time to start moving and to settle after the COM stops, so the bead transit time becomes longer than the molecular crossing time. Since the experiment measures only the bead, we compare the simulated bead transit time with the measured one. The time resolution is set by the macroscopic time step ΔT of one PCTP cycle. For a barrier region of a few nanometers crossed in tens of microseconds, as in DNA hairpin folding[46], a ΔT of a few microseconds (a few tens of the 200 ns internal steps) may resolve individual transit paths, and also keeps the COM excursion within the ∼1 nm criterion of Remark 1. Comparison with measured transit times will be the subject of our next study. It should show whether the brief pauses along a transition path[46] come from molecular intermediates or from bead and handle dynamics.

## 3. Conclusions

In summary, we have introduced force-probe coupled steered molecular dynamics (fp-SMD), a simulation framework that substantially narrows the long-standing timescale gap between all-atom SMD and optical tweezers SMFS by explicitly and self-consistently coupling the atomistic dynamics of a biomolecule to the physical dynamics of the force probe used to manipulate it. The accompanying Predict–Correct Trajectory Propagation (PCTP) scheme resolves the resulting computational bottleneck by alternating a bead-propagation step, with the molecular center of mass held fixed, and a molecular step in which metadynamics and transition-state theory with rate-matrix propagation identify the kinetically resolved intermediate reached at each macroscopic time step. On the cholesterol–β-CD host–guest complex, fp-SMD reproduces the experimental rupture-force histogram directly, without the systematic ∼4-fold overestimation characteristic of standard SMD, and, more significantly, directly resolves force-dependent switching of the molecular escape pathway, offering a structural basis for phenomena such as catch-slip bond behavior that have previously been accessible only indirectly. These results suggest that fp-SMD/PCTP can serve as a general framework for simulating force-probe-coupled biomolecular mechanics at experimentally relevant timescales, with a clear path toward larger and more complex mechanobiological systems.

## Supporting information

Revised Supporting Information

## ACKNOWLEDGMENTS

We thank our experimental collaborator and co-author of our previous work (ref. 6), Hanbin Mao at Kent State University, for valuable discussions on single-molecule force spectroscopy interpretation. This work was partially supported by the National Science Foundation (Grant No. 1953171). We are also grateful for the computing resources provided by the George Washington University (GW) Pegasus HPC facilities and NSF ACCESS facilities.

