## Supplementary material for "Simulating Bead-Tethered Biomolecules: Force-Probe Coupled Steered Molecular Dynamics (fp-SMD) in Optical Tweezers Force Spectroscopy": Revised Supporting Information

### S1. Validation of Bead Motion in a Harmonic Optical Trap

Before using the apparatus/bead parameters in Sections S2–S3, we first validate that the Predict–Correct Trajectory Propagation (PCTP) scheme in fp-SMD can correctly describe the Brownian motion of the bead and reproduce well-established, independently measured physics: the thermal motion of a bead in a harmonic optical trap, as directly measured by Florin and coworkers using optical trapping interferometry[1]. Figure S1 reproduces the mean-squared displacement (MSD) $\langle x^2(t)\rangle$ in their Fig. 1 (open symbols, digitized) for the two trap stiffnesses they report, $k_1$ = 1.2 µN/m and $k_3$ = 16 µN/m (bead mass $M = 8.5 \times 10^{-15}$ kg and radius a = 1.25 µm), overlaid with PCTP results run at the same stiffnesses and bead mass (solid lines). We also add two additional stiffer cases relevant to our previously published work[2]: the first one is k = 132.4 µN/m, bead mass $M = 5.04 \times 10^{-15}$ kg, and radius a = 1.05 µm –– the same trap stiffness and bead size used in our previous work[2], for which no experimental analog exists here; the second case uses the same a = 1.05 µm bead but fully tethered through the DNA linker ($k_{DNA} = 650$ µN/m) to cholesterol, with the cholesterol center of mass (COM) held fixed. This isolates the bead-trap-tether system from any molecular compliance and tests how the trap and DNA-tether stiffness combination sets the bead's dynamics (part i below), which is especially useful for estimating the maximum PCTP pulling distance per cycle in the linear-response regime (Section S2). We use the Gear backward differentiation formula to integrate the bead dynamics, with a timestep varying from 0.2 – 5 µs depending on the trap stiffness. For the three free-bead conditions (k1, k3, and k = 132.4 µN/m), the bead motion is only constrained by the harmonic optical trap, i.e., no tethered molecule/DNA linker attached to it, in contrast to the most rigid condition in the fourth case. Since no molecular relaxation is involved in MSD calculations, only PCTP Step 1 trajectory propagation (see the main text) is carried out.

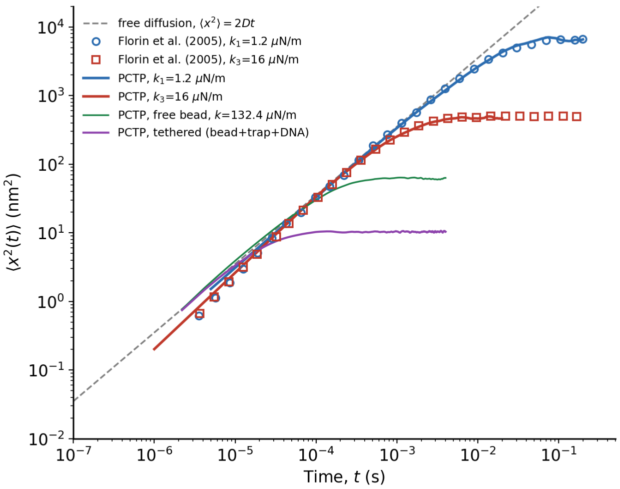


**Figure S1.** Mean-squared displacement (MSD) $\langle x^2(t) \rangle$ of a trapped bead vs. time, on log–log axes. Open circles/squares: experimental data of Florin et al. (2005) for $k_1$ = 1.2 µN/m and $k_3$ = 16 µN/m (bead mass $M = 8.5 \times 10^{-15}$ kg, radius a = 1.25 µm). Solid blue/red lines: PCTP results at the same two stiffnesses, showing close agreement with experiment across five decades of time. Solid green line: new PCTP result at k = 132.4 µN/m (bead mass $M = 5.04 \times 10^{-15}$ kg, radius a = 1.05 µm). Dashed gray line: free-diffusion reference $\langle x^2 \rangle = 2Dt$ using $D = k_BT / (6\pi\eta a)$ for the bead with radius a in water (η = 0.001 Pa·s) ($D \approx 1.76\times10^5$ nm²/s), which all curves approach at short times. Solid purple line: the same a = 1.05 µm bead fully tethered through the DNA linker to cholesterol (cholesterol center of mass (COM) held fixed), showing a much lower plateau (≈10 nm²) reflecting the added tether stiffness (part i).

**(i) Diffusion coefficient**. At short times ($t \ll \tau_A$) the bead has not yet felt the trap's restoring force, and all curves — experiment, PCTP, and the free-diffusion line — coincide, following $\langle x^2(t) \rangle = 2Dt$ with $D = k_BT/(6\pi\eta a)$, the Stokes–Einstein value for a free bead. This holds regardless of what is attached to the bead: expanding the exact MSD for a harmonic well, $MSD(t) = 2(k_BT/\kappa)(1 - e-t/\tau A)$, to first order in $t/\tau_A$ gives $MSD(t) \approx 2Dt$ — independent of κ. Three free-bead conditions appear in Fig. S1: the $k_1/k_3$ validation runs use a = 1.25 µm (matching Florin et al.'s largest sphere), giving $D \approx 1.76\times10^5$ nm²/s; the k = 132.4 µN/m run uses a smaller bead, a = 1.05 µm, giving $D \approx 2.09\times10^5$ nm²/s. The fourth curve uses the same a = 1.05 µm bead, now fully tethered through the DNA linker to cholesterol with cholesterol's COM held fixed. Because the trap and the DNA tether pull the bead from opposite sides against a fixed point (fixed cholesterol COM), they act as parallel springs on the same coordinate, $U(x) = ½k_{trap}(x-x_{trap})^2 + ½k_{DNA}(x-x_{chol,fixed})^2$, giving a combined stiffness $k_{eff} = k_{trap} + k_{DNA} = 132.4 + 650 = 782.4$ µN/m. Interestingly, this high stiffness is confirmed both by the fitted MSD plateau (10.29 ± 0.004 nm², yielding $k = 2k_BT/\text{plateau} = 804.9$ µN/m) and an independent equilibrium free-energy curvature $G''(q_U) = 0.0008$ N/m = 800 µN/m (Section S3, Fig. S6). Because the trap and DNA tether act in parallel directly on the bead, rather than in series through a compliant intermediate point, the resulting dynamics remain those of a single free bead relaxing in this stiffer combined well — governed by the bead's own Stokes–Einstein diffusion coefficient, not some artificially reduced effective value. This lets us extract D directly from the well's relaxation time and plateau: the $\tau_A$ fit for this curve (24.7 ± 0.2 µs) combined with $k_{eff}$ = 782.4 µN/m implies D ≈ 2.14×10$^5$ nm²/s, matching the free-bead value (2.09×10$^5$ nm²/s) within 2.4%. Based on this agreement, we use $D_q$ = 2.09×10$^5$ nm²/s as the bead's free diffusion coefficient throughout Sections S2–S3.

**(ii) Position autocorrelation (relaxation) time**. The trap's restoring force pulls a displaced bead back toward the trap center on a characteristic timescale $\tau_A = 6\pi\eta a/k = k_BT/(D\cdot k)$. Using the free-bead D ≈ 1.76×10$^5$ nm²/s established in part (i) for the a = 1.25 µm bead (matching Florin et al.'s own bead exactly, since the $k_1/k_3$ runs are untethered): $\tau_A(k_1)$ ≈ 19.6 ms and $\tau_A(k_3)$ ≈ 1.47 ms — both reasonably close to the values Florin et al. [1] report directly (≈20 ms and ≈1 ms, respectively), confirming PCTP reproduces not just the plateau amplitude ($2k_BT/k$) but the correct crossover timescale as well. For the stiffer free-bead condition, k = 132.4 µN/m, the plateau matches essentially exactly (fit = 62.9 ± 0.02 nm² vs. predicted $2k_BT/k$ ≈ 61.7 nm²), and the relaxation time $\tau_A$ = 154.1 ± 0.3 µs matches the prediction $\tau_A$ ≈ 149.5 µs using that run's own bead radius a = 1.05 µm and free-bead D ≈ 2.09×10$^5$ nm²/s (part i). For the tethered curve (bead + trap + DNA, cholesterol fixed): plateau = 10.29 ± 0.004 nm² and $\tau_A$ = 24.7 ± 0.2 µs. Both match the parallel-spring prediction closely: using $k_{eff}$ = 782.4 µN/m and the free-bead D = 2.09×10$^5$ nm²/s, the predicted plateau is 10.59 nm² (97.2% agreement) and predicted $\tau_A$ is 25.3 µs (97.5% agreement) — as good as the original $k_1/k_3$ validation against Florin et al. This confirms that $D_q$ for this system is simply the bead's own free-bead diffusion coefficient, 2.09×10$^5$ nm²/s, which we use throughout Sections S2–S3.

iii) **The two limiting regimes**. For $t \gg \tau_A$, the restoring force has had time to repeatedly act, and the bead's position has fully sampled the confined equilibrium (Boltzmann) distribution set by the trap; $\langle x^2(t)\rangle$ saturates at the constant value $2k_BT/k$ (visible as the flattening of the blue and red curves above). For $t \ll \tau_A$, the restoring force(s) have not yet had time to act, so the bead's motion is indistinguishable from that of a freely diffusing particle, independent of its position within the well. Comparing curves that share the same bead ($k_1$ vs. $k_3$, both a = 1.25 µm), the small residual spread at short times is sampling/binning noise; the k = 132.4 µN/m and tethered curves sit on a visibly different, steeper free-diffusion asymptote because they use a different, smaller bead (a = 1.05 µm, part i). This is why all four curves in Fig. S1 approach a free-diffusion line at short times, but not necessarily the same one. Ballistic motion on the shortest timescale observed by Florin et al.[1] is not studied here.

### S2. Maximum PCTP Pulling Distance per Cycle: Linear-Response (Hookean) Regime

Each PCTP prediction step (Step 1, see the main text) advances the bead by a step ΔR′ using a linear expansion of the activation force, $F_a (q, R) + (\partial F_a/\partial R)\, \Delta R'$. This linearization — and the Langer/Kramers treatment used throughout this SI — is only valid while the bead-linker system remains in the harmonic (Hookean) regime: $G(q) \approx \frac{1}{2}\kappa q^2$, with no significant anharmonic corrections. Below we use the raw equilibrium bead trajectory at F = 0 pN (corresponding to the bead fully tethered through the DNA linker to cholesterol in Fig. S1) to establish how far this harmonic approximation actually extends, and hence the largest per-cycle pulling distance ΔR for which quasi-equilibrium (linear-response) pulling is justified.

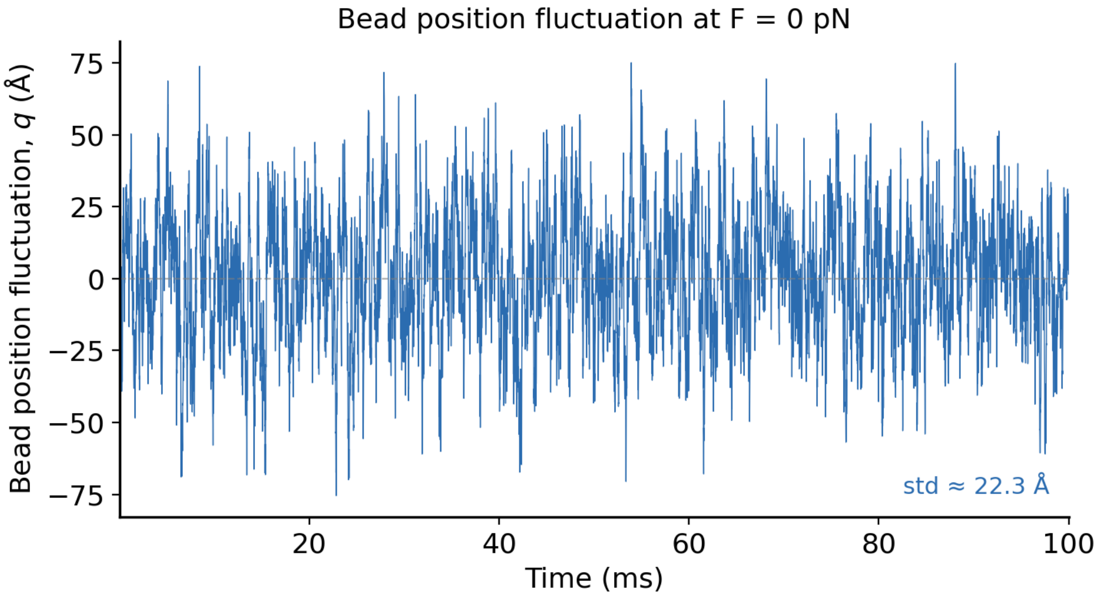


**Figure S2.** Bead position fluctuation q(t) at F = 0 pN, sampled every 0.01 ms over 100 ms (10,000 points). Mean ≈ 0.8 Å, standard deviation $\sigma_a \approx 22.3$ Å, range ≈ − 75 to +75 Å.

As shown in Fig. S2, the standard deviation provides an independent cross-check on the local curvature $G''(q_U)$ used below (Section S3): by equipartition for a harmonic well, $\sigma_a = \sqrt{(k_B T/\kappa)}$. Using $\kappa = G''(q_U) = 0.0008$ N/m (the force-resolved fit from Fig. S6) predicts $\sigma_a \approx 22.7$ Å — in close agreement with the ≈ 22.3 Å measured directly here. The governing criterion is the same Hookean-regime condition Cossio et al.[3] invoke for $k_{MA} \approx k_M$ to hold: $V''(q_U - x_U) < |G_0''(x_\cap)|$, i.e., the trap/linker stays softer than the molecular curvature and is not stretched into an anharmonic (near-contour-length) regime. Practically, this can be checked two ways: (i) statistically, from $\sigma_a$ via equipartition as above, and (ii) empirically, from where the measured free-energy profile $G(\zeta)$ (Fig. S6) departs from a clean parabola — which happens only beyond ±80–90 Å, where the sampling becomes sparse and noisy rather than reflecting real barrier structure.

**Formal bound on the safe pulling increment.** The discussion above establishes that *q* must stay within ±80–90 Å of its equilibrium position for the harmonic approximation to hold, but it does not involve how fast the bead can be driven through that region while still maintaining quasi-equilibrium. This requires treating the actual PCTP prediction step explicitly: over one macroscopic cycle of duration $\Delta T$, the driven coordinate advances by $\Delta R' \approx V \cdot \Delta T$, where V is the trap pulling velocity (R′ itself evolves continuously, but $\Delta R'$ is the net displacement accumulated over $\Delta T$). Two independent conditions must both hold for this step to qualify as quasi-equilibrium, linear-response pulling:

(a) Spatial condition: the step itself must not carry *q* outside the empirically validated Hookean region of G(q), namely, $\Delta R' \lesssim \Delta R_{max}$, with $\Delta R_{max}$ = 50 Å (≈ $2.2\sigma_a$, comfortably inside the ±80–90 Å quadratic region established from Fig. S6). This bounds the step size, independent of how it is reached in time.

(b) Temporal condition: the bead must have time to relax within its local harmonic well before the next step is taken. This means $\Delta T$ must be sufficiently long compared to the bead's own relaxation time, i.e., $\Delta T \gg \tau_A = k_BT/(D_q \cdot \kappa)$. Using $\kappa = G''(q_U) = 0.0008$ N/m — the force-resolved curvature established in Section S3, and $D_q = 2.09 \times 10^5$ nm²/s (the bead's own free-bead diffusion coefficient, Section S1), we have $\tau_A \approx 24.8$ µs.

Combining (a) and (b) through $\Delta R' = V \cdot \Delta T$ gives a single bound on the safe pulling velocity: since $\Delta T$ must satisfy both $\Delta T \gg \tau_A$ and $V \cdot \Delta T \leq \Delta R_{max}$, it follows that $V \ll \Delta R_{max} / \tau_A \approx 50$ Å / 24.8 µs ≈ $2.0\times10^5$ nm/s (≈ 202 µm/s).

This is the criterion Cossio et al.'s Hookean-regime condition[3] suggests for a driven (rather than purely equilibrium) trajectory: it is not enough for the linker to be Hookean in the static sense (V″ constant, established already); the pulling velocity must also stay well below $\Delta R_{max}/\tau_A$ so that $\Delta T$ can simultaneously satisfy $\Delta T \gtrsim \Delta R_{max}/V$ (to keep the step within $\Delta R_{max}$ at that speed) and $\Delta T \gg \tau_A$ (to stay quasi-equilibrium). In our optical-tweezers experiments and PCTP simulations, the pulling speed is restricted to V < 1000 nm/s (= 1 µm/s), which is at least 202 times below the derived bound of ≈202 µm/s — a comfortable safety margin confirming PCTP operates well within the quasi-equilibrium regime. With $\Delta R_{max} \approx 50$ Å satisfying condition (a), and $\tau_A \approx 24.8$ µs setting the timescale floor in condition (b), a per-cycle pulling distance of ΔR = 50 Å remains the recommended conservative choice.

### S3. Rate Pre-factor ($k_0$) Calculation for the K Matrix

To identify the kinetically resolved intermediate of cholesterol, the resulting master equation is propagated to the actual PCTP cycle time ($\Delta T$): $P(\Delta T) = \exp(-\mathbf{K}\,\Delta T)\,P(0)$, where the rate matrix **K** consists of all pairwise rate constants: $k_{ij} = k_0 * \exp(-\Delta G_{ij} / kT)$, where $\Delta G_{ij}$ is the local pair barrier between neighboring local minimum *i* and *j*, either forward or backward, and $k_0$ is a pre-factor which will be determined according to Cossio et al.[3]

Below we derive a single, state-independent $k_0$ from the molecule-linker-bead fluctuations, and apply it uniformly to every pairwise rate in **K**, with only the barrier $\Delta G$ varying by pair (from the metadynamics-derived free-energy surface, Fig. S7(C)).

Since the transition-state distance in Cossio et al.'s comparison of constant-force and constant-velocity pulling (their Eqs. 12–13) is on the nanometer scale[3], the resulting correction to the activation free energy is expected to fall below $k_B T$ for a linker as soft as our DNA handle ($k_{DNA} = 650$ μN/m). Consequently, we can treat the constant-force and constant-velocity rate constants as effectively interchangeable. We use Cossio et al.'s Eq. 16 (constant-force pulling) to relate the Langer rate $k_L$ to the molecular transition rate $k_M$

$$k_L = \frac{k_M\left[1-\gamma-\gamma\sigma+\sqrt{(1-\gamma-\gamma\sigma)^2+4\gamma\sigma}\right]}{2} \tag{1}$$

where $\sigma = D_q/D_x$ and $\gamma = V''(q_\cup - x_\cup)/|G''(x_\cap)|$.

To calculate the diffusion coefficients of the bead ($D_q$) and the cholesterol ($D_x$) in β-cyclodextrin (β-CD), we use two different integration strategies due to their distinct dynamics at very different timescales. For cholesterol, we fix the position of the bead because of its slow dynamics and perform a long molecular dynamics (MD) run to obtain the position distribution of cholesterol under thermal fluctuations. From the resulting trajectory, we compute the MSD (along the pulling direction) in the linear diffusive regime (Fig. S3) as a function of time and extract the diffusion coefficient via the Einstein relation: $\langle x^2(t)\rangle = 2Dt$. The diffusion coefficient of cholesterol is determined to be $D_x = 4.38 \times 10^8$ nm$^2$/s.

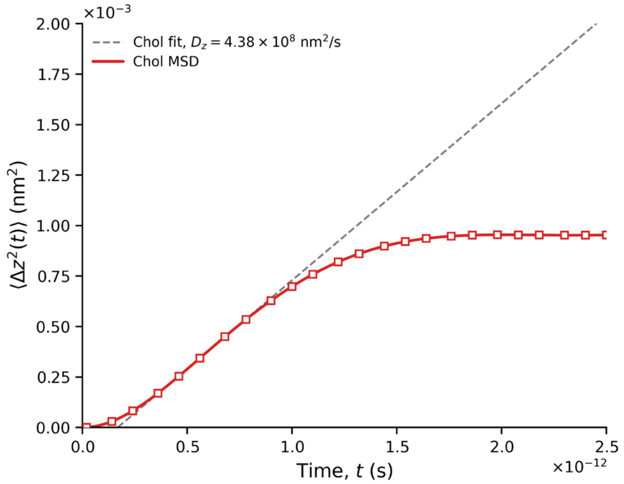

**Figure S3**. Mean-squared displacement (MSD) of the cholesterol center of mass (COM) versus time, from a fixed-bead MD trajectory. The linear fit (slope = $8.76 \times 10^8$ nm²/s) is used to extract the cholesterol diffusion coefficient $D_x$. The plateau at t ≈ 1 ps, $\tau_{mol}$, indicates the timescale over which cholesterol relaxes within its local confinement.

The bead diffusion coefficient $D_q$ is taken directly from the tethered-system analysis in Section S1 (Fig. S1): because the trap and DNA tether act on the bead from opposite sides against the fixed cholesterol, they combine as parallel springs ($k_{eff} = k_{trap} + k_{DNA}$, not in series), so the bead's dynamics reduce to a single free bead of its own radius diffusing in a stiffer combined well. This is confirmed by three independent, mutually consistent estimates of $k_{eff}$ (782–805 μN/m; see Section S1) and by the diffusion coefficient implied by the fit (≈$2.14\times10^5$ nm²/s) matching the free-bead value closely. We therefore use $D_q = 2.09\times10^5$ nm²/s throughout, giving a diffusion ratio $\sigma = D_q/D_x = 4.77\times10^{-4}$.

With the diffusion coefficients of both the bead and cholesterol determined, we further analyze the position fluctuations of the system to characterize the local free energy landscape. We accumulate the probability density $P(\zeta)$ as a function of the center-of-mass position $\zeta$ for both the bead and cholesterol (Fig. S4(a) for the bead, S4(b) for cholesterol), which reflects the system's fluctuations around the local minimum. The corresponding local free energy profile $G(\zeta)$ was then obtained via the Boltzmann relation:

$$G(\zeta) = -k_B T ln P(\zeta) \tag{2}$$

At the local minimum, the free energy was fitted to a quadratic form to extract the second derivative $G''(x_\cup)$ and $G''(q_\cup)$, which represents the local curvature. The fitted values are found to be $G''(x_\cup) = 6.28\ \mathrm{N/m}$ and $G''(q_\cup) = 0.0008\ \mathrm{N/m}$, see Figs. S5 and S6.

To calculate the ratio $\gamma$, we approximate the free energy curvature at the transition state $G''(x_\cap)$ by the curvature at the local minimum $G''(x_\cup)$. Furthermore, $V''(q_\cup - x_\cup)$ represents the linker spring stiffness, which is $6.5 \times 10^{-4}$ N/m. These yield $\gamma = 1.035 \times 10^{-4}$. Substituting $\sigma$ and $\gamma$ into Eq. (1), we find $k_L \approx k_M$.

Given that $k_L \approx k_M$ and $k_M \gg k_A$, the general relation below[3] (Eq. 3) simplifies to $k_{MA} \approx k_A$,

$$\frac{1}{k_{MA}} \approx \frac{1}{k_L} + \frac{1}{k_A} \tag{3}$$

indicating that the overall transition is primarily limited by the bead transition rate.

The bead transition rate $k_A$ can be estimated using one-dimensional Kramers theory along the bead coordinate $q$ (similar to Eq. 14 in Cossio et al. for calculating molecule transition rate[3])

$$k_A = \frac{\beta D_q}{2\pi}\sqrt{G''(q_{\cup})|G''(q_{\cap})|}e^{-\beta\Delta G^{\ddagger}} = k_0 e^{-\beta\Delta G^{\ddagger}} \quad (4)$$

where $\beta = 1/k_B T$ and $D_q$ is the bead diffusion coefficient. The second derivatives $G''(q_{\cup})$ and $G''(q_{\cap})$ represent the local curvatures at the minimum and the saddle point, respectively. Because our sampling (Figs. S4(a) and S6) resolves the well but not a clear barrier, we approximate these two curvatures as equal rather than measuring the saddle directly — this is an assumption, since Kramers/Langer pre-factors can be sensitive to this ratio if the true saddle curvature turns out to be substantially different.

We next accumulate the probability density $P(q)$ as a function of the bead position $q$, see Fig. S4(a). The corresponding free energy profile $G(q)$ is obtained by equation (2). The free energy is fitted to a quadratic form to extract the second derivative $G''(q_{\cup})$, which is 0.0011 kcal / (mol $\text{Å}^2$) = 0.0008 N/m (Fig. S6). The pre-factor $k_0$ of the Kramers transition rate is finally calculated as: $k_0 = \frac{\beta D_q}{2\pi} G''(q_{\cup}) = 6424.7\ s^{-1}$.

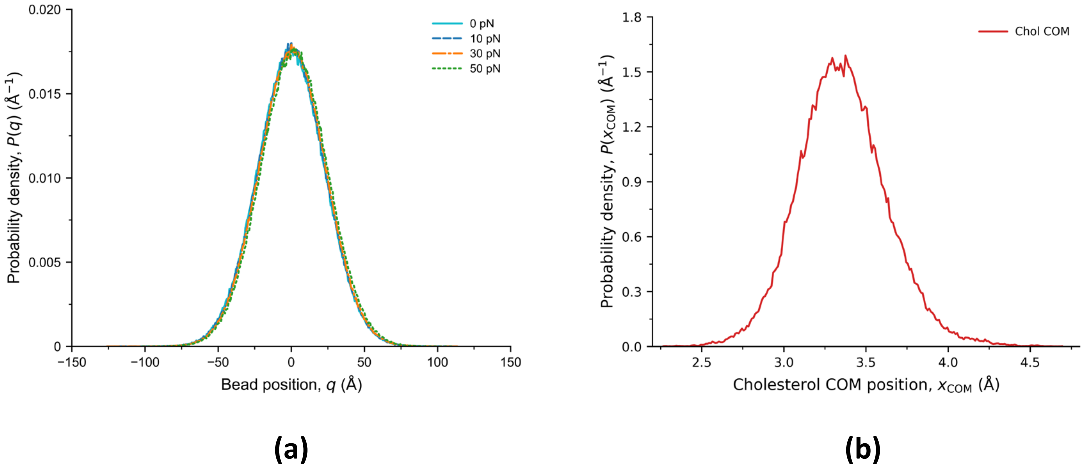


**Figure S4**. Probability densities of **(a)** P(q) of the bead position q, at applied forces of 0, 10, 30, and 50 pN; and **(b)** P($x_{com}$) of the cholesterol. The near-total overlap of the curves of P(q) indicates that the local well shape (and hence the curvature used below) is essentially force-independent over this range.

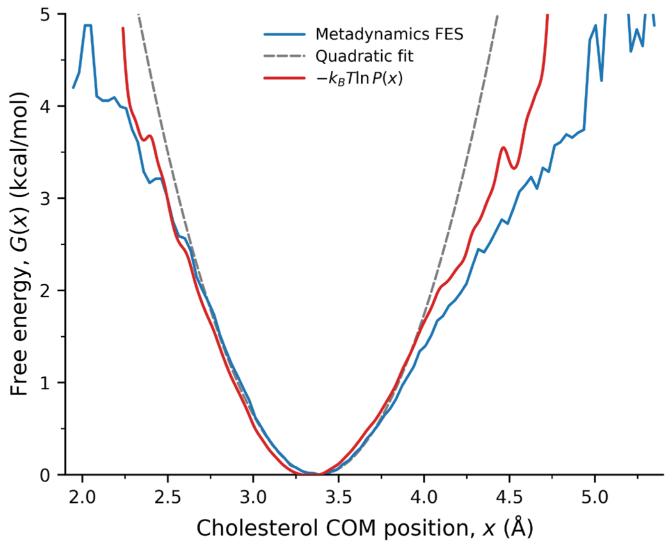


**Figure S5**. Free energy profiles for cholesterol along the pulling coordinate obtained from metadynamics FES (blue), from quadratic fit near the minimum (grey dashed), and from Fig. S4 (b) via Eq. (2) (red). All give G''($x_U$) ≈ 6.28 N/m for the γ calculation.

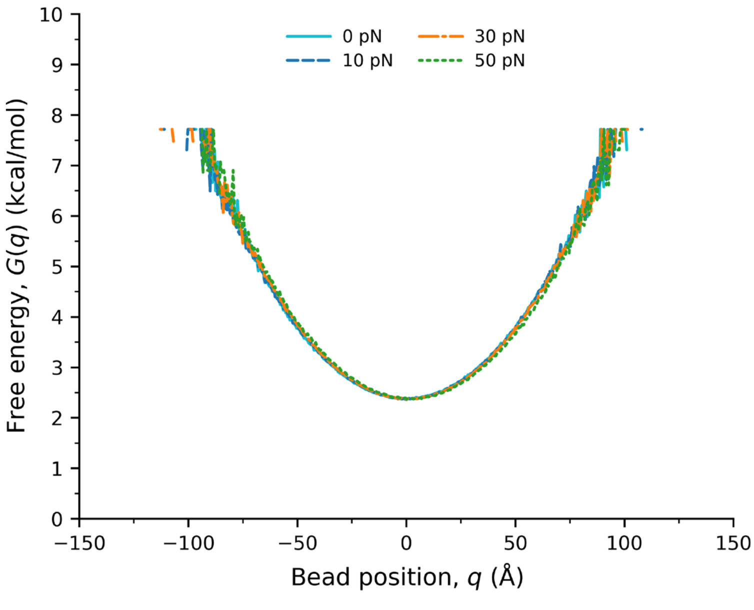


**Figure S6**. Free-energy profile $G(q) = -k_BT \ln P(q)$ along the pulling direction, obtained from Fig. S4 (a) via Eq. (2), at the same four forces. The quadratic fit near q = 0 gives the curvature G''($q_U$) — which is used in Eq. (4) for $k_0$ calculation. The noisy, sparsely sampled tails beyond ±80–90 Å cannot resolve any actual barrier/saddle point.

### S4. Mapping Free-Energy Surface (FES) and Potential of Mean Force (PMF) Profile of Cholesterol along Pulling Pathway under Different Loads

In the benchmark test of fp-SMD/PCTP simulation (see main text), the PCTP Step 2 molecular relaxation using well-tempered metadynamics sampling is the costliest step in one PCTP cycle, because mapping the free-energy surface (FES) of cholesterol in β-CD along the pulling pathway takes hundreds to thousands of Gaussians (equivalent to nanoseconds of regular MD) for cholesterol to fully explore the energy landscape of the host–guest complex. To overcome this difficulty, we pre-calculate all FESs — including the potential of mean force (PMF) profiles (see below) — at different loads and save them in a large dataset. This dataset also includes densely sampled meta-conformations of the host–guest complex, saved every few tens of Gaussians during metadynamics sampling. Because these meta-conformations still carry the artificial bias of the deposited Gaussians, they are de-biased using short (~2 ps) unbiased MD runs to obtain the actual kinetically resolved intermediates used in Step 2. In other words, pre-calculating and storing the FES and PMF profiles converts the most expensive step in every PCTP cycle into a one-time upfront cost, while the short de-biasing run in Step 2 is largely the only cost incurred per PCTP cycle.

In more detail, under each load we carry out a total of 20 metadynamics runs to improve the statistics. We set the load increment to 5 pN, spanning 0 to 65 pN — the maximum rupture force reported in the optical-tweezers experiment[2]. Here, we use the cholesterol center-of-mass position ($q_{com}$) and its tilt angle ($\theta$) away from the pulling direction as the two collective variables (CVs) used to map the two-dimensional (2D) FES. The tilt angle is particularly important for identifying the escaping pathway, given the large swing in $\theta$ near the exit region of β-CD. Figure S7(A) shows the evolution of the 2D FES with the external force applied to cholesterol, with the exit-side local minimum shifting gradually from a high-angle escape route to a low-angle route as the load increases.

Our metadynamics sampling also outputs a one-dimensional (1D) PMF profile along the pulling pathway — using only the cholesterol center-of-mass position ($q_{com}$) as the CV (Fig. S7(B)). The 1D PMF, averaged over the 20 metadynamics runs at each load, is used directly in propagating the master equation: $P(\Delta T) = \exp(-\mathbf{K}\Delta T)P(0)$. Figure S7(C) shows that the exit-side barrier for cholesterol decreases as the load increases — a well-known consequence of applying force along a reaction coordinate. In Kramers/Langer theory, as used throughout this SI, the force-dependent activation free energy is $\Delta G^{\ddagger}(F) = \Delta G^{\ddagger}(0) - F\cdot x^{\ddagger}$, where $x^{\ddagger}$ is the distance from the free-energy minimum to the transition state along the pulling direction (the same definition underlying $\Delta G^{\ddagger}_{M}$ in Cossio et al.'s[3] Eq. 14, Section S3 above). Applying force does mechanical work as cholesterol moves toward the exit, which tilts the free-energy landscape and lowers the barrier it must cross — the same physical picture behind the classic Bell–Evans description of force-accelerated bond rupture[4-6]. This is also the direct physical basis for the force-dependent Kramers rate, $k(F) = k_0 \cdot \exp(-\Delta G^{\ddagger}(F)/k_B T)$, that enters the pairwise rate constants of the **K** matrix (Section S3): as the load increases, $\Delta G^{\ddagger}(F)$ decreases, so the exit rate increases — exactly the trend seen in Fig. S7(C), in which the transition-state distance $x^{\ddagger}$ is about 10 Å over a large load range.

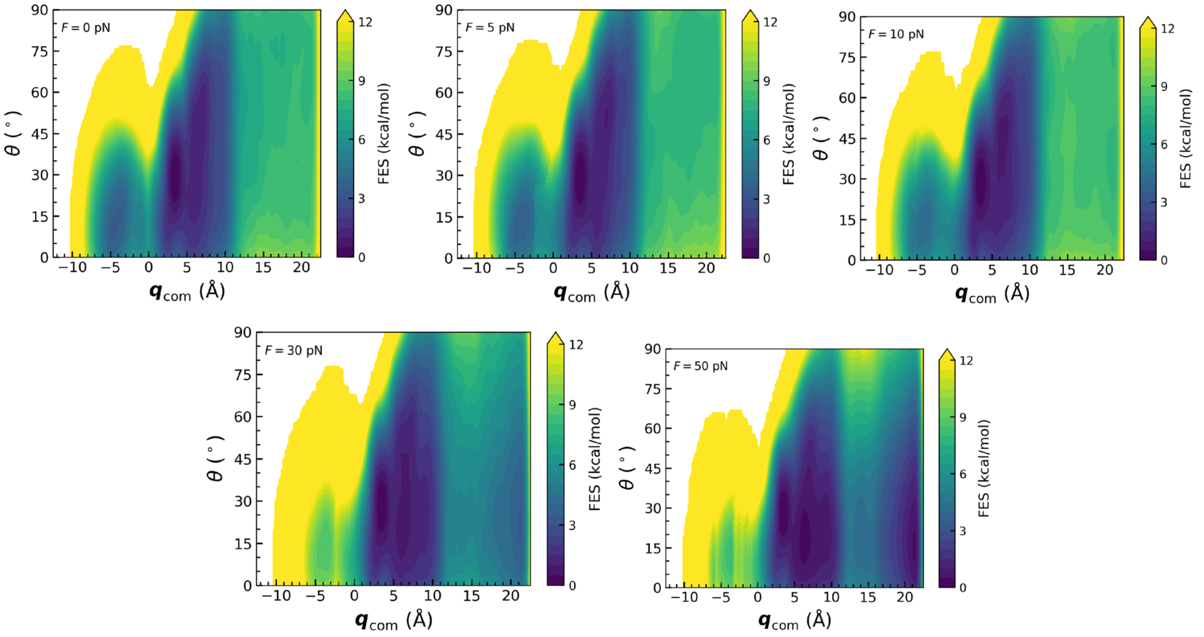


**A**

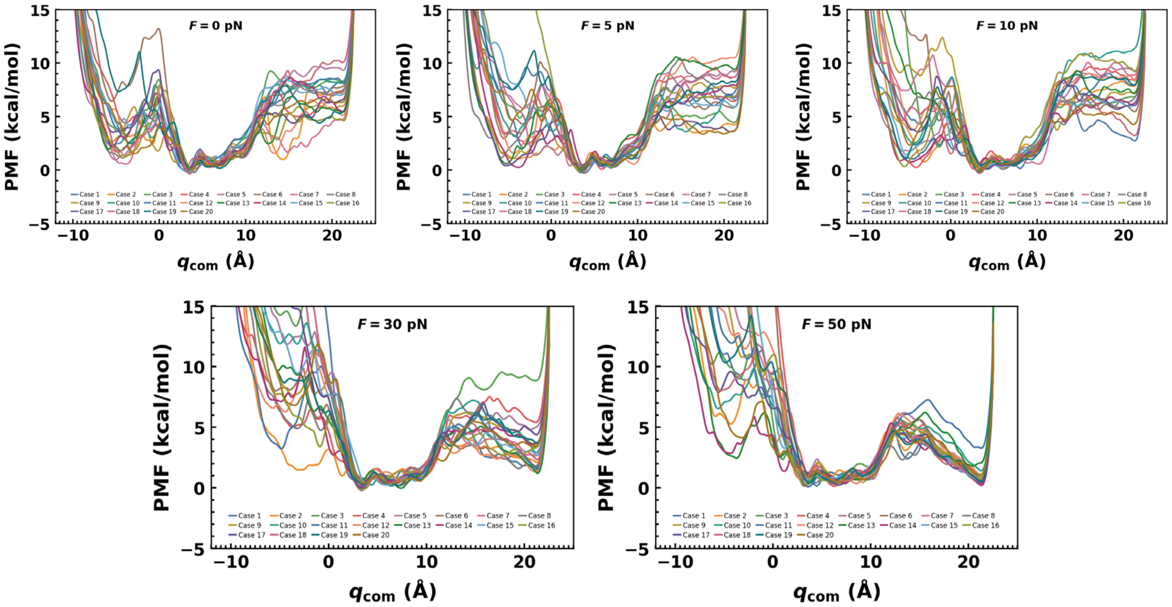


**B**

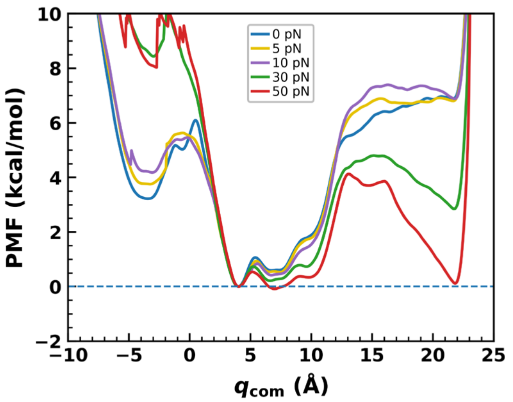


**C**

**Figure S7.** Free-energy surfaces (FESs) and potential-of-mean-force (PMF) profiles for cholesterol along the pulling pathway, pre-calculated during PCTP Step 2 metadynamics sampling at loads from 0 to 65 pN. **(A)** Two-dimensional (2D) FES, averaged over the 20 metadynamics runs per load, as a function of the cholesterol center-of-mass position ($q_{com}$) along the pulling direction and its tilt angle (θ) away from the pulling direction. **(B)** One-dimensional (1D) PMF profiles along $q_{com}$ from the individual metadynamics runs (up to 20 per load). **(C)** 1D PMF profiles averaged over the 20 runs per load. All three panels are shown for the same five loads (0, 5, 10, 30, and 50 pN). The transition-state distance $x^{‡}$ (the distance from the free-energy minimum to the transition state along the pulling direction) is about 10 Å over this load range. The exit-side barrier (positive $q_{com}$) decreases systematically with increasing load, consistent with the force-dependent Kramers rate discussed above.

### S5. Using Free-Energy Surfaces (FESs) and Potential of Mean Force (PMF) Profiles in fp-SMD/PCTP Simulation Runs

In each PCTP cycle, the cholesterol center-of-mass position ($q_{com}$) and tilt angle ($\theta$) carried over from the previous cycle are compared against those of the meta-conformations (all 20 independent replicates) stored for the new load predicted by the current PCTP Step 1 trajectory prediction. The meta-conformation with the closest-matching $q_{com}$ and $\theta$ is taken as the initial conformation for the 1D PMF profile used in the current cycle. After propagating the master equation (Section S3), a new $q_{com}$ is obtained; the meta-conformation whose $\theta$ most closely pairs with this new $q_{com}$ in the dataset is then identified as the new cholesterol meta-conformation. Following a short unbiased MD run (PCTP Step 2 molecular relaxation), the resulting $q_{com}$ and $\theta$ — which together uniquely define the new kinetically resolved intermediate of cholesterol, along with the corresponding β-CD conformation — are passed to PCTP Step 3 (trajectory correction). These updated $q_{com}$ and $\theta$ then serve as the initial conformation for the next PCTP cycle.

When performing PCTP simulation runs, we select either the lower-bound or upper-bound FES/PMF dataset to determine the kinetically resolved intermediate of cholesterol. The choice depends on which terminal force boundary—spanning 5n to 5(n+1) pN (where n = 0–12, see Section S4 in which the load increment is set to 5 pN, spanning 0 to 65 pN) —is closer to the actual force, $K_{trap}(Vt - R)$, in the current PCTP prediction step. Doing so is essentially controlling the "force" magnitude during the force-ramp pulling at constant velocity *V*, while the bead's actual displacement in each PCTP cycle time (ΔT) fluctuates around the average value V·ΔT.

We also test the displacement-controlled protocol to enforce the target per-cycle displacement (10 Å or 50 Å) exactly, letting the force fluctuate freely around its nominal ramp value. Doing so leads to a force fluctuation as large as ~1 pN, causing the bead to move between adjacent force-boundary datasets more readily. Because the Kramers escape rate depends exponentially on force, $k(F) = k_0 \cdot \exp(-\Delta G^{\ddagger}(F)/k_B T)$, and this rate is a convex function of F: an upward force excursion increases the escape rate more than an equal-sized downward excursion decreases it, so the two do not cancel on average. Averaged over a fluctuating force trajectory, this asymmetry raises the effective escape rate above what the trajectory's mean force alone would predict. The larger force fluctuations in the displacement-controlled protocol therefore increase the average escape rate at any given point in the ramp, causing rupture to occur earlier, i.e., at a systematically lower force (~ 5-6 pN decrease) than the smoother, force-controlled protocol. We therefore adopt the force-controlled force-ramp pulling.

### S6. Comparison between Standard SMD and fp-SMD/PCTP

Table S1 summarizes the qualitative and quantitative differences between standard SMD and fp-SMD/PCTP across the capabilities relevant to this benchmark.

**Table S1.** Capability comparison between standard SMD and fp-SMD/PCTP.

| Capability | Standard SMD | fp-SMD / PCTP |
|---|---|---|
| Physical model | Continuous atomistic trajectory; external force/velocity applied directly to the molecule; no explicit force-probe representation | Coupled Hamiltonian explicitly modeling molecule *and* bead (mass, drag, thermal noise); physically complete representation of the experiment |

| Capability | Standard SMD | fp-SMD / PCTP |
|---|---|---|
| Trajectory continuity | Single continuous, consistent atomistic trajectory (real momentum/velocity carried forward every step) | Not continuous; equilibrium free-energy sampling (metadynamics) across cycles |
| Key approximations | Single pulling coordinate; minimal added approximation beyond the force field itself | Molecule COM held fixed during Prediction; COM excursions of $\gtrsim 10$ Å tolerated (force change < 0.3 pN; SI Section S7); CV selection required for metadynamics (risk of missing hidden degrees of freedom) |
| Timescale accessible | ns–µs; must use unphysically fast pulling to observe rupture within reach | Bridges to ms–s (experimental) timescale via macroscopic $\Delta T$ cycles; resolution depends on $\Delta T$ |
| Loading-rate realism | Typically, $10^8$–$10^{11}$ pN/s; 6–11 orders of magnitude faster than experiment | Matches experimental loading rate directly (e.g., 5.5 pN/s) |
| Rupture-force accuracy | Systematically overestimates (~4× too high in this system) | Matches experimental histogram directly |
| Intermediate/pathway detection | True intermediates distorted by pulling-speed artifacts | Resolves kinetically-resolved intermediates via metadynamics + transition-state theory (rate-matrix propagation) at physical force; reveals force-dependent pathway switching |
| Escape/dissociation pathway | Pathway distorted by pulling-speed artifact; fast pulling can force a single, non-physiological route | Reveals true force-dependent pathway (high-angle vs. small-angle escape) — inaccessible to SMD |

| Capability | Standard SMD | fp-SMD / PCTP |
|---|---|---|
| Bound-state lifetime | Not physically meaningful; measured at an artificial pulling speed | Directly computable via transition-state theory (rate-matrix propagation) at physically realistic force levels and cycle times |
| Catch-slip bond behavior | Cannot be resolved — requires accurate lifetime vs. force across the correct regime, which SMD's speed artifacts preclude | In principle resolvable, since it is built from correctly scaled lifetime and force data — not yet demonstrated as a full curve |
| Scalability to complex systems | Scales in cost only, not physical realism | Requires free-energy landscape roughness and CV-adequacy/convergence testing for larger aggregates |

### S7. Bead Response to a Molecular COM Displacement (Frozen-COM Approximation in PCTP Step 1)

In PCTP Step 1, the bead equation (Eq. 8 of the main text) is integrated over ΔT with the cholesterol COM held at its previous-cycle thermodynamic average, $q_{com}$:

$$M \ddot{R} = -k_{DNA}(R - q_{com}) + K_{trap}(Vt - R) + F_{fr} + F_{th} \quad \text{(S7.1)}$$

Because the bead's response time, $\tau_A \approx 24.8$ µs (Section S2), is much shorter than ΔT (20–100 ms), the bead reaches local mechanical equilibrium within each cycle. Its mean position then follows from setting the net force to zero (friction, thermal noise and inertia average out):

$$R_{eq} = (k_{DNA} \cdot q_{com} + K_{trap} \cdot Vt) / (k_{DNA} + K_{trap}) \quad \text{(S7.2)}$$

The bead sits at the stiffness-weighted average of its two anchors: the molecular COM, reached through the DNA linker ($k_{DNA}$ = 650 µN/m), and the trap center ($K_{trap}$ = 132.4 µN/m; Section S1). If the COM moves by $\delta q_{com}$ while the trap center is fixed, the bead follows by

$$\Delta R = [k_{DNA}/(k_{DNA} + K_{trap})] \cdot \delta q_{com} = 0.83\ \delta q_{com} \quad \text{(S7.3)}$$

Because the linker is about five times stiffer than the trap, the bead follows the COM almost fully (for $\delta q_{com}$ = 1 nm, $\Delta R \approx 0.83$ nm). The linker extension therefore changes by only $\delta q_{com} - \Delta R = [K_{trap}/(k_{DNA} + K_{trap})]\cdot\delta q_{com} = 0.17\ \delta q_{com}$, and the force transmitted to the molecule changes by

$$\delta F = k_{DNA}\cdot 0.17\ \delta q_{com} = k_{series}\cdot\delta q_{com} \qquad \text{(S7.4)}$$

Here $k_{series} = k_{DNA}K_{trap}/(k_{DNA} + K_{trap}) \approx 110\ \mu N/m \approx 0.11$ pN/nm is the series stiffness of the linker–trap pair, which also sets the loading rate on the free molecule in the main text. The effect on the molecular kinetics follows from the force-dependent barrier, $\Delta G^{\ddagger}(F) = \Delta G^{\ddagger}(0) - F\cdot x^{\ddagger}$ (Section S4), so the rate exponent changes by

$$\delta \ln k = \delta F\cdot x^{\ddagger} / (k_B T) \qquad \text{(S7.5)}$$

with $x^{\ddagger} \approx 10$ Å (Fig. S7(C)) and $k_B T \approx 4.1$ pN·nm (T = 296 K). Table S2 evaluates Eqs. (S7.3) – (S7.5) for COM displacements of increasing size.

**Table S2.** Bead shift, transmitted-force change and rate-exponent change for a COM displacement $\delta q_{com}$ ($k_{DNA}$ = 650 μN/m, $K_{trap}$ = 132.4 μN/m, $x^{\ddagger}$ = 1 nm, $k_B T$ = 4.1 pN·nm).

| $\delta q_{com}$ (nm) | Meaning | $\Delta R$ (nm) | $\delta F$ (pN) | $\delta \ln k$ |
|---|---|---|---|---|
| 0.03 | Thermal COM fluctuation within the bound basin (Figs. S3, S4(b)) | 0.02 | 0.003 | 0.1% |
| 1.0 | Transition-state distance $x^{\ddagger}$ (Fig. S7(C)); reference value in the main text | 0.83 | 0.11 | 3% |
| 1.8 | Full COM range of the PMF, primary minimum (≈ 4 Å) to exit plateau (≈ 22 Å) (Fig. S7(C)) | 1.5 | 0.20 | 5% |
| 8 | Upper end of the Hookean range of the linker (Section S2) | 6.6 | 0.88 | 21% |

Three consequences follow. (i) Thermal fluctuations of the COM within a basin (≈ 0.03 nm) occur on the picosecond timescale (Fig. S3), far faster than the bead's response time ($\tau_A \approx 24.7$ μs), so the bead responds only to the mean COM position, which is the frozen value used in Step 1. (ii) If a conformational transition during $\Delta T$ displaces the COM by more than $x^{\ddagger}$, Step 2 identifies the new state through the master equation, its COM is taken from the updated conformational ensemble, and Step 3 solves Eq. (S7.1) again with the new $q_{com}$. This removes the Step 1 error in the predicted bead position (Eq. S7.3), and because the molecular force changes by only ≈ 0.11 pN per nm of COM motion (Eq. S7.4), the Step 2–3 loop (convergence threshold ≈ 1 nm, main text) needs at most one additional pass. (iii) The validity of the frozen-COM approximation is therefore set by force, not by the molecular relaxation time: it holds as long as $k_{series} \cdot \delta q_{com}$ remains well below $k_B T/x^{\ddagger} \approx 4$ pN and well below the 5 pN width of the force bins used to select the FES/PMF datasets (Section S5). Within the Hookean range of the linker ($\delta q_{com} \lesssim 8$ nm), $\delta F < 1$ pN. Larger conformational rearrangements, stiffer linkers, or much longer $\Delta T$ would require smaller $\Delta T$ or additional Step 2–3 iterations, and mapping these regimes is left for future work.
